# Natural selection and circular pathways to seasonal migration in birds

**DOI:** 10.1101/2020.07.20.212217

**Authors:** Matthew R. Halley

## Abstract

The “migratory revolutions” (MR) model is a synthetic theory of bird migration that seeks to explain the full range of the functional phenotype, from sedentary residents of non-seasonal (tropical) habitats to obligate long-distance migrants, as a cumulative evolutionary response to shifting distributions of adult extrinsic mortality across the annual cycle. At macroevolutionary scales, the general model predicts that migration evolves in circular patterns, reframing classic debates about the effects of migration on speciation and extinction rates. Here, I describe and apply the MR model to a well-known system, the passerine genus *Catharus* (Turdidae), to illustrate its broad implications for reconstructing evolutionary history.

---

“All that we can do, is to keep steadily in mind that each organic being is striving to increase in a geometrical ratio; that each at some period of its life, during some season of the year, during each generation, or at intervals, has to struggle for life and to suffer great destruction.” (*1*)

The intertwined evolutionary causes and consequences of bird migration have fascinated and perplexed biologists for centuries (*1*–*5*). A recent wave of research has greatly expanded the discussion, shining light on the polygenic nature of the migratory phenotype (e.g. *6,7*), functional links between behavior and morphology (*8,9*), the sources and targets of selection (*10–16*), fundamental tradeoffs between fecundity and survival (*14*), and the heterobathmic distribution of behavioral and morphological components of the migratory phenotype among populations and species (*5,6,17*). That the expression of migratory behavior originates (in sedentary populations) as an evolutionary response to high mortality during the resource-depleted non-breeding season (*11*–*14*) is an alternative hypothesis that has recently gained traction over the traditional view of migration as an adaptation by which tropical species increase fecundity by exploiting seasonal (pulsed) food resources in temperate regions (*18,19*). However, a general macroevolutionary theory that incorporates this mechanism has remained elusive.

Here, I advance a synthetic theory of the evolutionary sequence(s) that produced the observed spectrum of migratory behavior in birds. The “migratory revolutions” (MR) model conceptually integrates morphology and behavior (*20,21*) and presumes that extreme optimization for long-distance migration (*14,22,23*) constrains the directionality of evolutionary change. The model explains (predicts) the complete spectrum of the migratory phenotype, from sedentary residents of non-seasonal habitats, to obligate long-distance migrants that also practice short-distance (“intratropical”) migration during the non-breeding season (*24,25*). To my knowledge, no other general model of the evolution of bird migration accomplishes these core objectives.

Migratory behavior and its functional morphology are polygenic traits with extensive phenotypic integration, that presumably evolve in tandem as a single “functional phenotype” (*20,21*). Embryonic growth rates do not differ between migratory and non-migratory taxa (*14*), and forelimbs (wings, W) and hindlimbs (legs, L) are ontogenetically independent and achieve adult size within the first month after hatching. Notwithstanding this independence, there is an inverse relationship between W and L investment during development, presumably because of performance tradeoffs, which produces, at macroevolutionary scales, a negative relationship between wing length and tarsometatarsus length (*26*). Hyper-aerial and migratory species have the highest W/L ratios; selection presumably favors longer wings and/or shorter legs, which reduce weight and aerodynamic drag (*27*). Flightless and otherwise sedentary birds have the lowest W/L ratios; after colonizing islands with few predators, lineages predictably evolve “downstream” toward flightlessness (low W/L ratio), after accounting for island size and phylogenetic history (*28*). Evolution in the “upstream” direction on the W/L axis, at least during the transition from sedentary behavior to short-distance migration, is driven by an asymmetrical annual distribution of adult extrinsic mortality (AEM), *i*.*e*., higher mortality during the non-breeding (winter) period in temperate regions (*11*–*13*).

In the MR model, the functional phenotype evolves upstream in response to the iterative and predictable development of “point sources” of selection (periodic high-mortality events) that are asymmetrically distributed across the annual cycle (Fig. 1). Point sources are best viewed as a predictable (annual) series of filters through which a subset of individuals pass. The model depends on the assumption that a quantitative trait evolving via directional selection will not reverse its (continuous) evolutionary trajectory unless the source of selection is diminished or removed, or a stronger source of selection is applied in the opposite direction. Therefore, some point sources are “persistent” (predictable annual events) and constrain evolutionary reversal. Short-distance migration evolves in response to a persistent selective force: mortality during the non-breeding season, which is proximately caused by resource depletion and/or interspecific competition (*11*–*13*). Long-distance migration evolves from short-distance migration in response to a different, persistent selective force: additive migration-related mortality (e.g. crossing the Caribbean Sea during the Atlantic hurricane season, *16*), which optimizes the functional phenotype for annual adult survival at the cost of annual fecundity (*14*). In the model, long-distance migrants can only “escape” this positive feedback loop by forming colonies. If colonization occurs on an oceanic island, the lineage will evolve toward low W/L ratio and the MR cycle will stop (*28*). Whereas, in continental settings (e.g. *31,32*), newly sedentary colonies may expand toward temperate regions, reinitiating the cycle.

**Figure 1.**
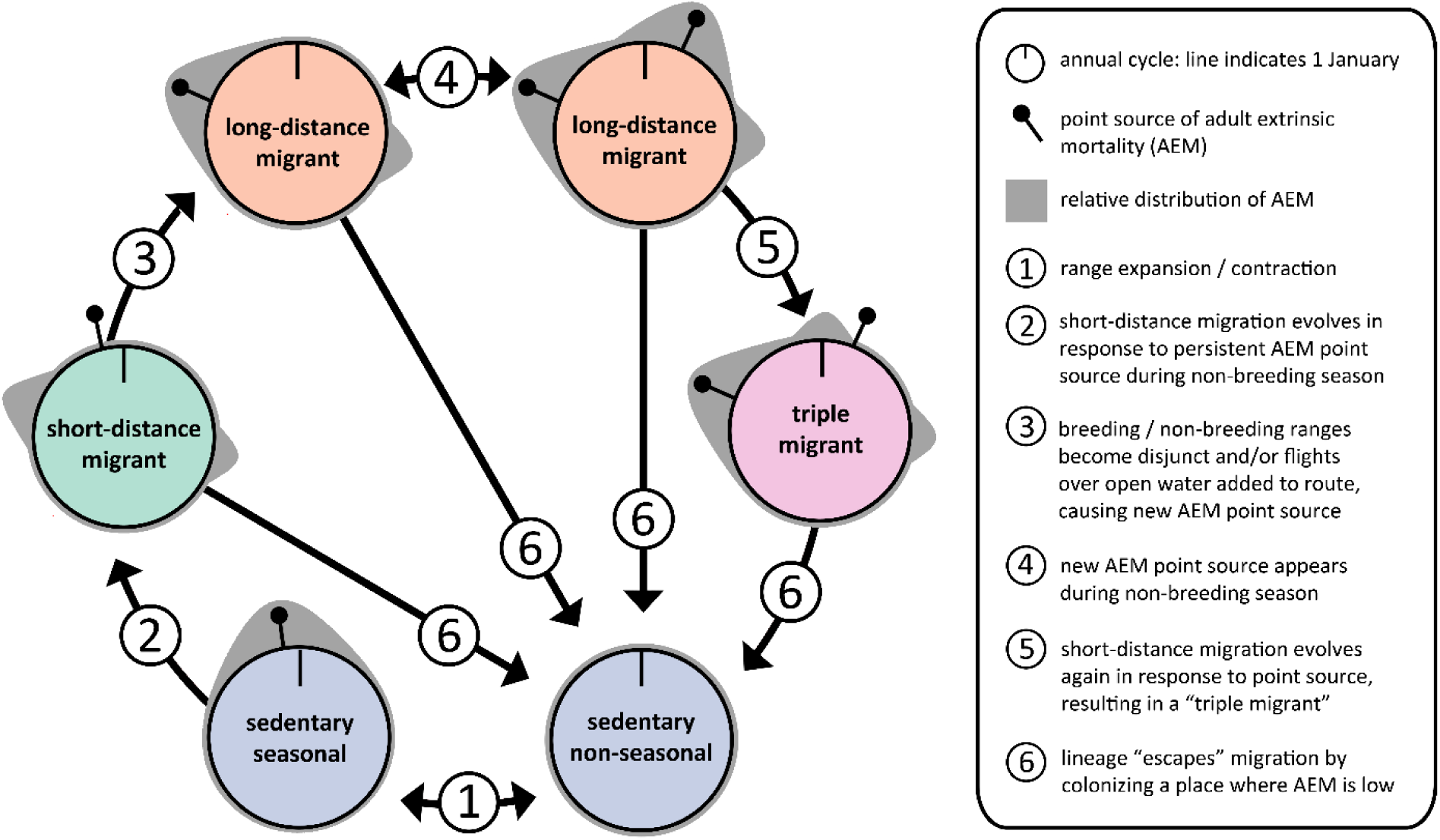
The migratory revolutions (MR) model of the evolution of avian migration. Each circle represents the annual cycle of a typical Nearctic-Neotropical migrant. Time proceeds in a clockwise direction. By rotating various elements, the model can be adjusted to accommodate taxa in other geographical contexts (e.g. migration within the southern hemisphere). Gray shading shows the relative distribution of adult extrinsic mortality (AEM) across the year, not absolute differences among migratory states. For sedentary species in non-seasonal environments (bottom right, purple), predation is the main source of AEM, which is lower and more evenly distributed across the annual cycle than in temperate-breeding species (*29,30*). A sedentary species that expands into temperate (seasonal) regions (bottom left, purple) is subjected to new (additive) sources of AEM during the non-breeding (winter) period (*11*–*13*), which emerge as a point source of selection (black pin), disproportionately eliminating sedentary individuals and driving the evolution of short-distance migration (left, green). Subsequently, the breeding and non-breeding ranges of short-distance migrants may become geographically disjunct, and/or long-distance flights over open water may be incorporated into the migratory route. This produces a new persistent source of selection (high AEM) caused by migration-related mortality during the first migration after breeding (i.e. fall migration), driving the evolution of long-distance migration (top left, orange) (*14,31*). The subsequent emergence of yet another source of AEM during the non-breeding season of a long-distance migrant (top right, orange) may cause short-distance migration to evolve a second time, resulting in triple migration (right, pink). Evolutionary reversal is precluded after the migratory phenotype is gained and “escape” from the highly optimized migratory phenotype is only possible by forming a sedentary colony.

To demonstrate how the MR model generates novel hypotheses about evolutionary history, consider a well-known model system, the genus *Catharus* (Aves: Turdidae), which exhibits the gamut of migratory behavior including a short-distance migrant (*C*. *guttatus*), a “triple migrant” (*C*. *fuscescens*), and numerous sedentary and long-distance migrants (e.g. *C*. *gracilirostris* and *C*. *minimus*, respectively). *Catharus* forms a species-rich crown group that is flanked by a paraphyletic stem group comprised of a monotypic long-distance migrant, *Hylocichla mustelina* (sister to *Catharus*), and several monotypic and depauperate genera that are mostly sedentary (*33*). Phylogenetic evidence indicates that the common ancestor of extant *Catharus* species diverged approximately 5 mya, and the common ancestor of *Catharus* and *H*. *mustelina* diverged ≤ 8 mya (*33*). Remarkably, migration is homoplastic: *C*. *guttatus*, the only short-distance migrant, and the only species in *Catharus* that remains in temperate North America in winter, is more closely related to three sedentary species than to the long-distance migratory species which it superficially resembles (*33*–*36*).

Previous attempts to reconstruct the ancestral state of migration in *Catharus* used a binary coding scheme (migratory/sedentary) and grouped *C*. *guttatus* with the other migrants (*33,34*). However, the MR model predicts that (1) short-distance migrants have intermediate morphology and behavior, and (2) short-distance migration may be a more stable evolutionary state, because AEM is more evenly distributed across the annual cycle (i.e. no point sources, see Fig. 1) and short-distance migrants do not pay the cost of lower annual fecundity (*14*). As predicted, the short-distant migrant *C*. *guttatus* has intermediate W/L morphology in both sexes, relative to congeners with sedentary (lower W/L) and long-distance and triple migrant (higher W/L) phenotypes (Fig. 2). Nevertheless, the intermediate morphology is still capable of sporadic feats of extreme dispersal, even across large water bodies, as evidenced by a hatch-year *C*. *guttatus* photographed in the Azores in October 2019 (*37*). This implies that colonization does not depend on long-distance migration evolving first (see Fig. 1, transition 6).

**Figure 2.**
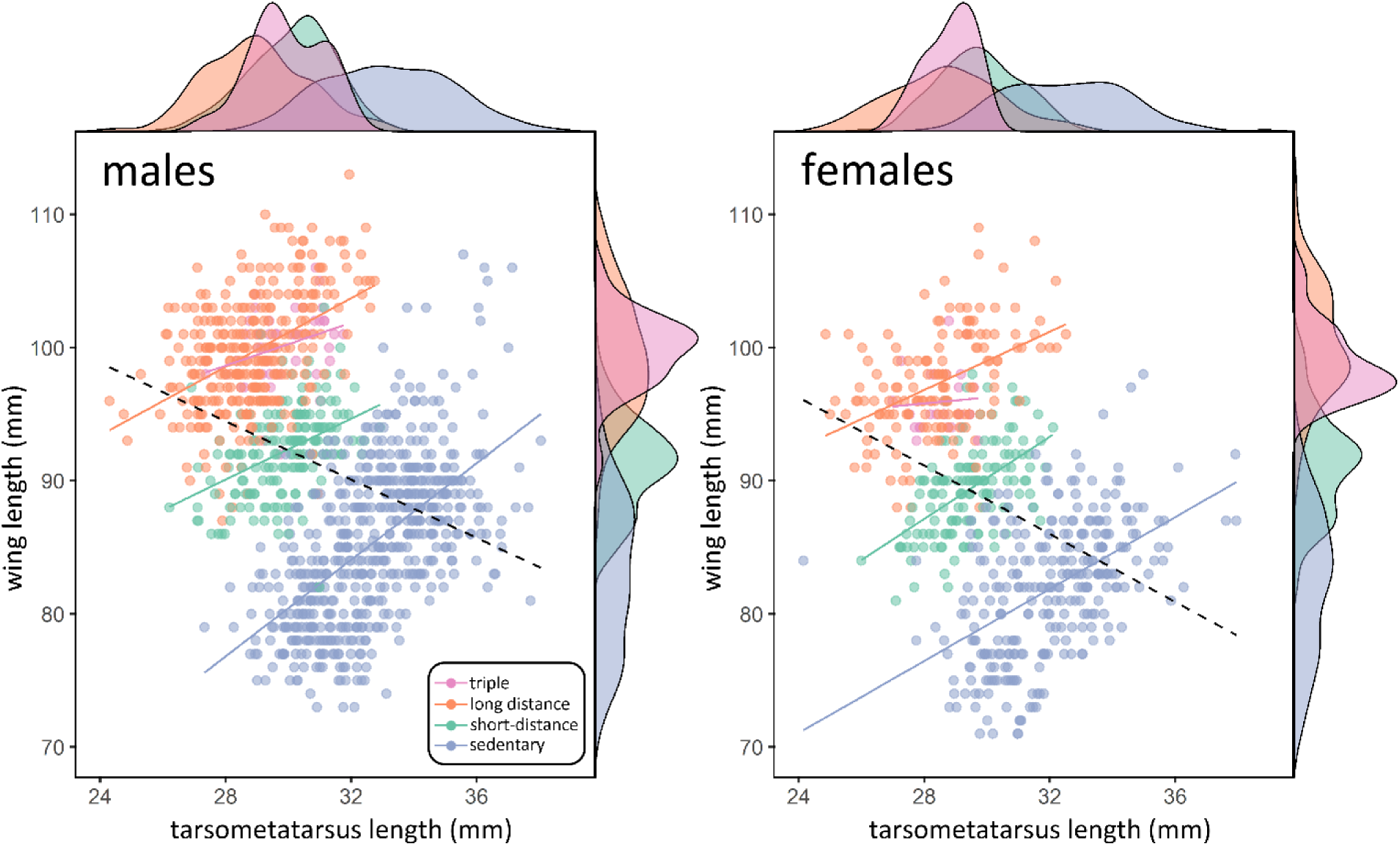
Functional morphology of migration in the genus *Catharus* in relation to sex. Separate plots are shown for males (*n* = 1,096) and females (*n* = 603). As predicted by the migratory revolutions (MR) model, the short-distance migrant (*C*. *guttatus*) occupies an intermediate (green) morphological space, between sedentary (purple) and long-distance and triple migratory states (orange, pink). Evolutionary shifts in the migratory phenotype occur in an “upstream” direction: linear regression lines (dashed black) of the complete male and female datasets, respectively, have significantly negative slopes (males: *R*^*2*^ = 0.11, *β* = −1.095, *p* ≪ 0.001; females: *R*^*2*^ = 0.14, *β* = −1.279, *p* ≪ 0.001). In contrast, within each migratory class (except female triple migrants), regression lines (colored) have significantly positive slopes (see Supplementary Materials). Density plots are shown for both axes. Color scheme is standardized across figures.

By mapping MR model transitions to the *Catharus* phylogeny, we can evaluate the most parsimonious forms of three hypotheses concerning the ancestral state of migration (Fig. 3). (1) The “sedentary ancestor hypothesis” posits that the migratory phenotype evolved three times independently, with no recent losses of migration. (2) The “short-distance migratory ancestor hypothesis” posits that *C*. *guttatus* is a lineage in evolutionary stasis (i.e. long-lived with a stable phenotype). This novel hypothesis explains the apparent homoplasy of the migratory plumage phenotype without invoking convergent evolution (i.e., it is actually homology, *contra 35*), and explains the ecological incumbency of *C*. *guttatus* in North America during winter (which other hypotheses ignore). It also implies that the evolution of long-distance migration in the common ancestor of the long-distance clade may have been a response to competitive exclusion with its (incumbent) parental lineage (*C*. *guttatus*) during winter. Indeed, ecological niche modeling suggests that *C*. *guttatus*, alone among the migratory *Catharus* species, likely maintained a widespread breeding range in North America during glacial maxima, and therefore may have been relatively buffered from the influence of the Pleistocene climate cycles (*39*). The intermediate *C*. *guttatus* also maintains a “fast” life history, relative to long-distance migratory *Catharus* species that have the derived condition of lower annual fecundity (*14*). (3) The “long-distance migratory ancestor hypothesis” posits that, on at least two occasions, migratory lineages colonized the Neotropics and evolved low W/L ratios. Those colonies subsequently diversified to form the two major sedentary clades in *Catharus* (*33*). This hypothesis treats short-distance migration as an autapomorphic condition in *C*. *guttatus* and invokes convergent evolution to explain the similarity of its external phenotype to the long-distance migrants, although the hypothesized ecological mechanisms driving the convergence remain vague (*35*).

**Figure 3.**
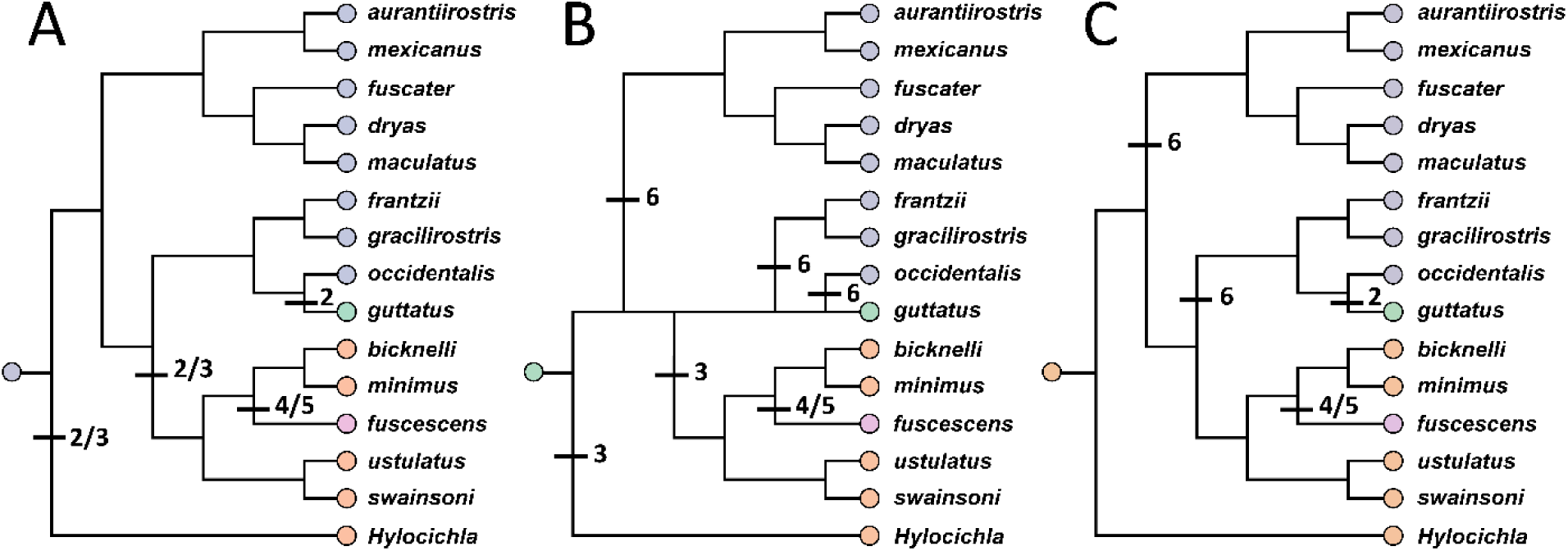
Phylogenetic hypotheses of the evolution of migration in the genus *Catharus*. Cladogram topology follows (*33*) and (*36*) with taxonomic updates by (*38*). Color scheme is standardized across figures. Numbered transitions between migratory states are standardized with Fig. 1. **(A)** The “sedentary ancestor hypothesis” infers that the migratory phenotype evolved three times independently. **(B)** The “short-distance migratory ancestor hypothesis” posits that the *C*. *guttatus* lineage is in evolutionary stasis (visualized here by straightening its branch) and infers three independent losses of migration (colonization events) and one gain of long-distance migration. **(C)** The “long-distance migratory ancestor hypothesis” infers that the migratory phenotype was lost twice (via colonization) and subsequently regained in *C*. *guttatus*.

All three ancestral state hypotheses explain the “intratropical migration” of *C*. *fuscescens* as an autapomorphic condition (i.e. triple migration), contrary to early speculations that it may represent an ancestral state (*25*, but see below about circularity). However, other evolutionary transitions in the MR cycle invoke different selection pressures to explain the observed phenotypes, and transitions among the various states probably occur at different rates, complicating the process of directly comparing and evaluating these hypotheses. Nevertheless, given the relative stability of the short-distance migratory phenotype in the MR model, and the fact that the intermediate (short-distance) hypothesis is capable of explaining the totality of the observed data (including the ecological incumbency of *C*. *guttatus* in winter), the intermediate (short-distance) hypothesis should be considered the working (null) hypothesis for now.

Lastly, a novel prediction of the MR model is that the functional phenotype evolves in a circular pattern at macroevolutionary scales. Multiple “migratory revolutions” can theoretically occur in a single lineage, and the number of revolutions is predicted to increase with time. This fundamentally reframes the long-standing debate about the effects of the migratory phenotype on speciation and extinction rates, which ultimately shape patterns of biodiversity (*1,2,40,41*). Unlike previous suggestions that lineages “switch” between migratory and sedentary states (*39*), the MR model requires that sedentary colonies (descended from long-distance migrants) pass through the short-distance migratory phenotype before the long-distance phenotype is regained (i.e. evolutionary change is continuous, see Fig. 2). If the MR model is true, we have no method to ascertain the average period of the long-term cycle, and it is unclear whether the model can be applied beyond its first iteration. We can only presume that (currently) sedentary groups that are descended from migratory ancestors (that were themselves descended from sedentary ancestors, etc.) have retained much of the functional genetic variation necessary for short-distance migration to be re-expressed, when or if they expand again into temperate regions (*6,21*). Therefore, we would not expect to find sequence divergence at many migration-related genes in comparisons of closely related sedentary and migratory taxa (e.g. *8,42*). If evolutionary pathways to seasonal migration are indeed circular, as predicted by the MR model, it is probable that the original assembly of the genetic variation underlying the spectrum of the functional phenotype was an extremely ancient event, perhaps antecedent to the origin of modern birds (*21,43*).

## Acknowledgments

I thank Jason D. Weckstein, Christopher M. Heckscher, Mark B. Robbins, Therese A. Catanach, Jeremy J. Kirchman, and the late Amotz Zahavi, for enlightening conversations that inspired this work. Nathan H. Rice, Paul Sweet, Chris M. Milensky, and Jean L. Woods provided access to specimen collections. Funding for this project came from ANSP, Drexel University, and the William L. McLean III Fellowship awarded to MRH.

## References

1. C. Darwin, On the Origin of Species (John Murray, London, UK, 1859).

2. T. H. Montgomery, Jr., Extensive migration in birds as a check upon the production of geographical varieties. Am. Nat. 30, 458–464 (1896).

3. G. H. Lowery, Jr., Evidence of trans-Gulf migration. Auk 63, 175–211 (1946).

4. D. J. Levey, F. G. Stiles, Evolutionary precursors of long-distance migration: resource availability and movement patterns in Neotropical landbirds. Am. Nat. 140, 447–476 (1992).

5. R. M. Zink, Towards a framework for understanding the evolution of avian migration. J. Avian Biol. 33, 433–436 (2002).

6. R. M. Zink, The evolution of avian migration. Biol. J. Linn. Soc. 104, 237–250 (2011).

7. K. E. Delmore, D. P. L. Toews, R. R. Germain, G. L. Owens, D. E. Irwin, The genetics of seasonal migration and plumage color. Curr. Biol. 26, 2167–2173 (2016).

8. M. Lundberg, M. Liedvogel, K. Larson, H. Sigeman, M. Grahn, A. Wright, S. Åkesson, S. Bensch, Genetic differences between willow warbler migratory phenotypes are few and cluster in large haplotype blocks. Evol. Lett. 1, 155–168 (2017).

9. M. S. Bowlin, Sex, Wingtip Shape, and Wing-Loading Predict Arrival Date at a Stopover Site in the Swainson’s Thrush (*Catharus ustulatus*). Auk 124, 1388–1396 (2007).

10. C. Sheard, M. H. C. Neate-Clegg, N. Alioravainen, S. E. I. Jones, C. Vincent, H. E. A. MacGregor, T. P. Bregman, S. Claramunt, J. A. Tobias, Ecological drivers of global gradients in avian dispersal inferred from wing morphology. Nat. Commun. 11, 2463 (2019).

11. V. Salewski, B. Bruderer, The evolution of bird migration—a synthesis. Naturwissenschaften 94, 268–279 (2007).

12. A. K. Shaw, Drivers of animal migration and implications in changing environments. Evol. Ecol. 30, 991–1007 (2016).

13. B. M. Winger, G. G. Auteri, T. M. Pegan, B. C. Weeks, A long winter for the Red Queen: rethinking the evolution of seasonal migration. Biol. Rev. 94, 737–752 (2019).

14. B. M. Winger, T. M. Pegan, https://doi.org/10.1101/2020.06.27.175539 (2020).

15. T. S. Sillett, R. T. Holmes, Variation in survivorship of a migratory songbird throughout its annual cycle. J. Anim. Ecol. 71, 296–308 (2002).

16. C. M. Heckscher, A Nearctic-Neotropical migratory songbird’s nesting phenology and clutch size are predictors of accumulated cyclone energy. Sci. Rep. 8, 9899 (2018).

17. B. Helm, E. Gwinner, Migratory restlessness in an equatorial nonmigratory bird. PLoS Biology 4, e110. https://doi.org/10.1371/journal.pbio.0040110

18. G. Cox, The role of competition in the evolution of migration. Evolution 22, 180–192 (1968).

19. G. Cox, The evolution of avian migration systems between the temperate and tropical regions of the New World. Am. Nat. 126, 451–474 (1985).

20. R. C. Bertossa, Morphology and behaviour: functional links in development and evolution. Philos. T. R. Soc. B. 366, 2056–2068 (2011).

21. H. Dingle, Animal migration: is there a common migratory syndrome? J. Ornithol. 147, 212–220 (2006).

22. D. Sol, N. Garcia, A. Iwaniuk, K. Davis, A. Meade, W. A. Boyle, T. Szekely, Evolutionary divergence in brain size between migratory and resident birds. PLoS ONE 5, 1–8.

23. D. L. DeAngelis, W. M. Post, C. C. Travis, Positive feedback in natural systems (Springer-Verlag, New York, NY, 1986).

24. B. A. Loiselle, J. G. Blake, Temporal variation in birds and fruits along an elevational gradient in Costa Rica. Ecology 72, 180–193 (1991).

25. C. M. Heckscher, M. R. Halley, P. Stampul, Intratropical migration of a Nearctic-Neotropical migratory songbird (*Catharus fuscescens*) in South America with implications for migration theory. J. Trop. Ecol. 31, 285–289 (2015).

26. A. M. Heers, K. P. Dial, Wings versus legs in the avian *bauplan*: Development and evolution of alternative locomotor strategies. Evolution 69, 305–320 (2014).

27. C. J. Pennycuick, Modeling the flying bird. Volume 5. (Academic Press, Cambridge, MA, 2008).

28. N. A. Wright, D. W. Steadman, C. C. Witt, Predictable evolution toward flightlessness in volant island birds. Proc. Natl. Acad. Sci. 201522931 (2016)

29. T. E. Martin, Avian life-history evolution has an eminent past: does it have a bright future? Auk 121, 289–301 (2004).

30. T. E. Martin, M. M. Riordan, R. Repin, J. C. Mouton, W. M. Black, Apparent annual survival estimates of tropical songbirds better reflect life history variation when based on intensive field methods. Glob. Ecol. Biogeogr. 12, 1386–1397 (2017).

31. B. M. Winger, F. K. Barker, R. H. Ree, Temperate origins of long-distance seasonal migration in New World songbirds. Proc. Natl. Acad. Sci. 111, 12115–12120 (2014).

32. V. Gómez-Bahamón, R. Marquez, A. E. Jahn, C. Yumi Miyaki, D. T. Tuero, O. Laverde-R., S. Restrepo, C. D. Cadena, Speciation associated with shifts in migratory behavior in an avian radiation. Curr. Biol. 30, 1312–1321 (2020).

33. G. Voelker, R. C. K. Bowie, J. Klicka, Gene trees, species trees and Earth history combine to shed light on the evolution of migration in a model avian system. Mol. Ecol. 22, 3333–3344 (2013).

34. D. C. Outlaw, G. Voelker, B. Mila, D. J. Girman, Evolution of long-distance migration in and historical biogeography of *Catharus* thrushes: a molecular phylogenetic approach. Auk 120, 299–310 (2003).

35. K. Winker, C. L. Pruett, Seasonal migration, speciation, and morphological convergence in the genus *Catharus* (Turdidae). Auk 123, 1052–1068 (2006).

36. K. M. Everson, J. F. McLaughlin, I. A. Cato, M. M. Evans, A. R. Gastaldi, K. K. Mills, K. G. Shink, S. M. Wilbur, K. Winker, Speciation, gene flow, and seasonal migration in *Catharus* thrushes (Aves: Turdidae). Mol. Phylo. Evol. 139, 106564 (2019).

37. Macaulay Library, Cornell University (ML182488001).

38. M. R. Halley, The misidentification of *Turdus ustulatus* Nuttall, and the names of the nightingale-thrushes (Turdidae: *Catharus*). Bull. Brit. Orn. Cl. 139, 248–69 (2019).

39. R. M. Zink, A. S. Gardner, Glaciation as a migratory switch. Sci. Adv. 3, e1603133 (2017).

40. K. Winker, On the origin of species through heteropatric differentiation: a review and a model of speciation in migratory animals. Ornithol. Monogr. 69, 1–30 (2010).

41. C. R. Marshall, Five palaeobiological laws needed to understand the evolution of the living biota. Nat. Ecol. Evol. 1, 0165 (2017).

42. J. S. Lugo Ramos, K. E. Delmore, M. Liedvogel, Candidate genes for migration do not distinguish migratory and non-migratory birds. J. Comp. Physiol. A 203, 383–397 (2017).

43. P. Berthold, A comprehensive theory for the evolution, control and adaptability of avian migration. Ostrich 70, 1–11 (1999).

