## Supplementary Materials for "Natural selection and circular pathways to seasonal migration in birds"

Matthew R. Halley[1,2*]

[1]Department of Biodiversity, Earth and Environmental Science, Drexel University, Philadelphia, Pennsylvania, USA.

[2]Ornithology Department, Academy of Natural Sciences of Drexel University, Philadelphia, Pennsylvania, USA.

---

### Materials and Methods

MRH recorded measurements from a large sample of adult study skins of known sex (1,096 males, 603 females) in the collections of the American Museum of Natural History (AMNH), Academy of Natural Sciences of Drexel University (ANSP), Delaware Museum of Natural History (DMNH), Field Museum of Natural History (FMNH), and National Museum of Natural History, Smithsonian Institution (USNM). The following data were taken: (1) wing length, equal to the flattened distance between the carpal joint and the tip of the longest primary remige, measured with a metric ruler, and (2) tarsometatarsus length, equal to the distance between the intertarsal joint and the distal end of the final leg scale, measured with calipers. Each study skin was assigned to a taxonomic group based on its external morphology and collection locality. The dataset was nearly taxonomically comprehensive at the subspecies level, containing >90% of named taxa in *Catharus* and all major species complexes (see list in Halley 2019, *Bull. Brit. Orn. Cl.* 139, 248–69). All statistical analyses were performed with RStudio (R Core Team, 2014) with code provided below. Morphometric data will be deposited in Dryad Repository after peer review.

### R Code used in analysis

```
CATHARUSDATA <- read.table("WG_TAR_KNOWNSEX.csv", header = TRUE, sep = ",");
CATHARUSDATA_M <- subset(CATHARUSDATA, SEX == "M")
CATHARUSDATA_M_R <- subset(CATHARUSDATA_M, MIGR3 == "R")
CATHARUSDATA_M_M <- subset(CATHARUSDATA_M, MIGR3 == "M")
CATHARUSDATA_M_G <- subset(CATHARUSDATA_M, MIGR3 == "G")
CATHARUSDATA_M_T <- subset(CATHARUSDATA_M, MIGR3 == "T")

CATHARUSDATA_F <- subset(CATHARUSDATA, SEX == "F")
CATHARUSDATA_F_R <- subset(CATHARUSDATA_F, MIGR3 == "R")
CATHARUSDATA_F_M <- subset(CATHARUSDATA_F, MIGR3 == "M")
CATHARUSDATA_F_G <- subset(CATHARUSDATA_F, MIGR3 == "G")
CATHARUSDATA_F_T <- subset(CATHARUSDATA_F, MIGR3 == "T")
```

```
####LinearModels####
####CompleteDataset####
linreg_ALL <- lm(WL~TAR, data = CATHARUSDATA)
summary(linreg_ALL)
confint(linreg_ALL)

####FemalesOnly####
linreg_F <- lm(WL~TAR, data = CATHARUSDATA_F)
summary(linreg_F)
confint(linreg_F)

linreg_F_G <- lm(WL~TAR, data = CATHARUSDATA_F_G)
summary(linreg_F_G)
confint(linreg_F_G)

linreg_F_M <- lm(WL~TAR, data = CATHARUSDATA_F_M)
summary(linreg_F_M)
confint(linreg_F_M)

linreg_F_R <- lm(WL~TAR, data = CATHARUSDATA_F_R)
summary(linreg_F_R)
confint(linreg_F_R)

linreg_F_T <- lm(WL~TAR, data = CATHARUSDATA_F_T)
summary(linreg_F_T)
confint(linreg_F_T)

####MalesOnly####
linreg_M <- lm(WL~TAR, data = CATHARUSDATA_M)
summary(linreg_M)
confint(linreg_M)

linreg_M_G <- lm(WL~TAR, data = CATHARUSDATA_M_G)
summary(linreg_M_G)
confint(linreg_M_G)

linreg_M_M <- lm(WL~TAR, data = CATHARUSDATA_M_M)
summary(linreg_M_M)
confint(linreg_M_M)

linreg_M_R <- lm(WL~TAR, data = CATHARUSDATA_M_R)
summary(linreg_M_R)
confint(linreg_M_R)

linreg_M_T <- lm(WL~TAR, data = CATHARUSDATA_M_T)
summary(linreg_M_T)
confint(linreg_M_T)
```

### Statistical Results (RStudio output)

```
> ####CompleteDataset####
> linreg_ALL <- lm(WL~TAR, data = CATHARUSDATA)
> summary(linreg_ALL)

Call:
lm(formula = WL ~ TAR, data = CATHARUSDATA)

Residuals:
    Min      1Q  Median      3Q     Max 
-20.144  -5.167   0.720   5.024  24.142 

Coefficients:
             Estimate Std. Error t value Pr(>|t|)    
(Intercept) 122.50979    2.43847   50.24   <2e-16 ***
TAR          -1.05360    0.07889  -13.36   <2e-16 ***
---
Signif. codes:  0 ‘***’ 0.001 ‘**’ 0.01 ‘*’ 0.05 ‘.’ 0.1 ‘ ’ 1

Residual standard error: 7.789 on 1697 degrees of freedom
Multiple R-squared:  0.09512,	Adjusted R-squared:  0.09459 
F-statistic: 178.4 on 1 and 1697 DF,  p-value: < 2.2e-16

> confint(linreg_ALL)
                 2.5 %      97.5 %
(Intercept) 117.727052 127.2925201
TAR          -1.208327  -0.8988811

> ####FemalesOnly####
> linreg_F <- lm(WL~TAR, data = CATHARUSDATA_F)
> summary(linreg_F)

Call:
lm(formula = WL ~ TAR, data = CATHARUSDATA_F)

Residuals:
     Min       1Q   Median       3Q      Max 
-17.8671  -4.5131   0.4293   4.7800  21.3718 

Coefficients:
            Estimate Std. Error t value Pr(>|t|)    
(Intercept)  126.953      3.962  32.047   <2e-16 ***
TAR           -1.279      0.130  -9.841   <2e-16 ***
---
Signif. codes:  0 ‘***’ 0.001 ‘**’ 0.01 ‘*’ 0.05 ‘.’ 0.1 ‘ ’ 1
```

```
Residual standard error: 7.296 on 601 degrees of freedom
Multiple R-squared:  0.1388,	Adjusted R-squared:  0.1373 
F-statistic: 96.85 on 1 and 601 DF,  p-value: < 2.2e-16

> confint(linreg_F)
                 2.5 %     97.5 %
(Intercept) 119.172839 134.732785
TAR          -1.534643  -1.024023
> linreg_F_G <- lm(WL~TAR, data = CATHARUSDATA_F_G)
> summary(linreg_F_G)

Call:
lm(formula = WL ~ TAR, data = CATHARUSDATA_F_G)

Residuals:
    Min      1Q  Median      3Q     Max 
-6.2865 -2.0394 -0.0299  1.8011  8.5105 

Coefficients:
            Estimate Std. Error t value Pr(>|t|)    
(Intercept)  43.9471     6.0574   7.255 2.70e-11 ***
TAR           1.5422     0.2056   7.500 7.25e-12 ***
---
Signif. codes:  0 ‘***’ 0.001 ‘**’ 0.01 ‘*’ 0.05 ‘.’ 0.1 ‘ ’ 1

Residual standard error: 2.878 on 137 degrees of freedom
Multiple R-squared:  0.2911,	Adjusted R-squared:  0.2859 
F-statistic: 56.24 on 1 and 137 DF,  p-value: 7.253e-12

> confint(linreg_F_G)
                2.5 %    97.5 %
(Intercept) 31.969030 55.925096
TAR          1.135598  1.948886
> linreg_F_M <- lm(WL~TAR, data = CATHARUSDATA_F_M)
> summary(linreg_F_M)

Call:
lm(formula = WL ~ TAR, data = CATHARUSDATA_F_M)

Residuals:
    Min      1Q  Median      3Q     Max 
-10.739  -2.430  -0.222   2.196  10.294 

Coefficients:
            Estimate Std. Error t value Pr(>|t|)
```

```
(Intercept)  66.2114     5.1417   12.88  < 2e-16 ***
TAR          1.0930     0.1804    6.06 1.08e-08 ***
---
Signif. codes:  0 ‘***’ 0.001 ‘**’ 0.01 ‘*’ 0.05 ‘.’ 0.1 ‘ ’ 1

Residual standard error: 3.541 on 148 degrees of freedom
Multiple R-squared:  0.1988,	Adjusted R-squared:  0.1934
F-statistic: 36.72 on 1 and 148 DF,  p-value: 1.078e-08

> confint(linreg_F_M)
                2.5 %    97.5 %
(Intercept) 56.0508359 76.371920
TAR          0.7365615  1.449399
> linreg_F_R <- lm(WL~TAR, data = CATHARUSDATA_F_R)
> summary(linreg_F_R)

Call:
lm(formula = WL ~ TAR, data = CATHARUSDATA_F_R)

Residuals:
   Min     1Q Median     3Q    Max
-9.502 -3.232 -0.082  2.796 12.732

Coefficients:
            Estimate Std. Error t value Pr(>|t|)
(Intercept)  38.6652     4.4634   8.663  3.5e-16 ***
TAR          1.3500     0.1391   9.703  < 2e-16 ***
---
Signif. codes:  0 ‘***’ 0.001 ‘**’ 0.01 ‘*’ 0.05 ‘.’ 0.1 ‘ ’ 1

Residual standard error: 4.428 on 285 degrees of freedom
Multiple R-squared:  0.2483,	Adjusted R-squared:  0.2457
F-statistic: 94.15 on 1 and 285 DF,  p-value: < 2.2e-16

> confint(linreg_F_R)
               2.5 %    97.5 %
(Intercept) 29.879793 47.450644
TAR          1.076136  1.623853
> linreg_F_T <- lm(WL~TAR, data = CATHARUSDATA_F_T)
> summary(linreg_F_T)

Call:
lm(formula = WL ~ TAR, data = CATHARUSDATA_F_T)

Residuals:
   Min     1Q Median     3Q    Max
```

```
-3.182 -1.745 -0.883  1.450  6.029

Coefficients:
            Estimate Std. Error t value Pr(>|t|)
(Intercept)  89.5002    17.6596   5.068 3.13e-05 ***
TAR           0.2248     0.6177   0.364    0.719
---
Signif. codes:  0 ‘***’ 0.001 ‘**’ 0.01 ‘*’ 0.05 ‘.’ 0.1 ‘ ’ 1

Residual standard error: 2.342 on 25 degrees of freedom
Multiple R-squared:  0.005271,	Adjusted R-squared:  -0.03452
F-statistic: 0.1325 on 1 and 25 DF,  p-value: 0.7189

> confint(linreg_F_T)
               2.5 %     97.5 %
(Intercept) 53.12964 125.870764
TAR         -1.04730   1.496948

> ####MalesOnly####
> linreg_M <- lm(WL~TAR, data = CATHARUSDATA_M)
> summary(linreg_M)

Call:
lm(formula = WL ~ TAR, data = CATHARUSDATA_M)

Residuals:
     Min       1Q   Median       3Q      Max
-18.3692  -5.0097   0.8384   4.8677  22.8494

Coefficients:
             Estimate Std. Error t value Pr(>|t|)
(Intercept) 125.11659    3.00320   41.66   <2e-16 ***
TAR          -1.09474    0.09641  -11.36   <2e-16 ***
---
Signif. codes:  0 ‘***’ 0.001 ‘**’ 0.01 ‘*’ 0.05 ‘.’ 0.1 ‘ ’ 1

Residual standard error: 7.728 on 1094 degrees of freedom
Multiple R-squared:  0.1054,	Adjusted R-squared:  0.1046
F-statistic: 128.9 on 1 and 1094 DF,  p-value: < 2.2e-16

> confint(linreg_M)
                 2.5 %      97.5 %
(Intercept) 119.223898 131.0092816
TAR          -1.283907  -0.9055698
> linreg_M_G <- lm(WL~TAR, data = CATHARUSDATA_M_G)
> summary(linreg_M_G)
```

```
Call:
lm(formula = WL ~ TAR, data = CATHARUSDATA_M_G)

Residuals:
     Min       1Q   Median       3Q      Max
-11.4886  -1.9147   0.0856   1.9088   9.6355

Coefficients:
            Estimate Std. Error t value Pr(>|t|)
(Intercept)  58.1089     5.4482   10.67  < 2e-16 ***
TAR           1.1420     0.1821    6.27 2.82e-09 ***
---
Signif. codes:  0 ‘***’ 0.001 ‘**’ 0.01 ‘*’ 0.05 ‘.’ 0.1 ‘ ’ 1

Residual standard error: 2.997 on 172 degrees of freedom
Multiple R-squared:  0.1861,	Adjusted R-squared:  0.1813
F-statistic: 39.32 on 1 and 172 DF,  p-value: 2.815e-09

> confint(linreg_M_G)
                 2.5 %    97.5 %
(Intercept) 47.3549128 68.862897
TAR          0.7825193  1.501516
> linreg_M_M <- lm(WL~TAR, data = CATHARUSDATA_M_M)
> summary(linreg_M_M)

Call:
lm(formula = WL ~ TAR, data = CATHARUSDATA_M_M)

Residuals:
    Min      1Q  Median      3Q     Max
-13.343  -2.668   0.003   2.626   9.819

Coefficients:
            Estimate Std. Error t value Pr(>|t|)
(Intercept)  62.5299     4.2723  14.636   <2e-16 ***
TAR           1.2868     0.1479   8.703   <2e-16 ***
---
Signif. codes:  0 ‘***’ 0.001 ‘**’ 0.01 ‘*’ 0.05 ‘.’ 0.1 ‘ ’ 1

Residual standard error: 3.933 on 303 degrees of freedom
Multiple R-squared:    0.2,	Adjusted R-squared:  0.1973
F-statistic: 75.74 on 1 and 303 DF,  p-value: < 2.2e-16

> confint(linreg_M_M)
                 2.5 %    97.5 %
```

```
(Intercept) 54.1228503 70.937003
TAR         0.9958149  1.577728
> linreg_M_R <- lm(WL~TAR, data = CATHARUSDATA_M_R)
> summary(linreg_M_R)

Call:
lm(formula = WL ~ TAR, data = CATHARUSDATA_M_R)

Residuals:
     Min       1Q   Median       3Q      Max
-13.4302  -3.3835  -0.3519   3.0884  16.4669

Coefficients:
            Estimate Std. Error t value Pr(>|t|)
(Intercept)  26.0577     3.4990   7.447 3.59e-13 ***
TAR           1.8126     0.1068  16.978  < 2e-16 ***
---
Signif. codes:  0 ‘***’ 0.001 ‘**’ 0.01 ‘*’ 0.05 ‘.’ 0.1 ‘ ’ 1

Residual standard error: 4.804 on 564 degrees of freedom
Multiple R-squared:  0.3382,	Adjusted R-squared:  0.337
F-statistic: 288.2 on 1 and 564 DF,  p-value: < 2.2e-16

> confint(linreg_M_R)
                2.5 %    97.5 %
(Intercept) 19.185035 32.930451
TAR          1.602928  2.022336
> linreg_M_T <- lm(WL~TAR, data = CATHARUSDATA_M_T)
> summary(linreg_M_T)

Call:
lm(formula = WL ~ TAR, data = CATHARUSDATA_M_T)

Residuals:
    Min      1Q  Median      3Q     Max
-8.0359 -1.3687  0.3847  1.1572  5.0549

Coefficients:
            Estimate Std. Error t value Pr(>|t|)
(Intercept)  76.1152     9.8643   7.716  5.2e-10 ***
TAR           0.8046     0.3313   2.429   0.0189 *
---
Signif. codes:  0 ‘***’ 0.001 ‘**’ 0.01 ‘*’ 0.05 ‘.’ 0.1 ‘ ’ 1

Residual standard error: 2.499 on 49 degrees of freedom
Multiple R-squared:  0.1075,	Adjusted R-squared:  0.08924
```

```
F-statistic: 5.899 on 1 and 49 DF,  p-value: 0.01886

> confint(linreg_M_T)
                2.5 %    97.5 %
(Intercept) 56.2921700 95.938199
TAR          0.1388884  1.470312
```
